# Exploring the virome of *Gyropsylla spegazziniana*, a major yerba mate pest

**DOI:** 10.64898/2026.04.01.715862

**Authors:** Yesica Gisel Candia, Vanesa Nahirñak, Alejandra Badaracco, Humberto Debat, María Elena Schapovaloff, Nicolás Bejerman

## Abstract

The yerba mate psyllid (*Gyropsylla spegazziniana*) poses a significant threat to yerba mate crops, causing extensive economic losses. While some ecological aspects as well as control strategies have been studied, its associated viral diversity remains unexplored. Here, by generating the first RNA high-throughput analysis (HTS) of this pest, we explored the *G. spegazziniana* virome, revealing novel and diverse RNA viruses. We characterized five new viral members belonging to distinct families, with evolutionary cues of beny-like viruses (*Benyviridae*), picorna-like viruses (*Picornaviridae*), and sobemo-like viruses (*Solemoviridae*); which were tentatively named Gyropsylla spegazziniana beny-like virus 1 (GSBlV1), Gyropsylla spegazziniana picorna-like virus 1 (GSPlV1), and Gyropsylla spegazziniana sobemo-like virus 1-3 (GSSlV1-3), respectively. Phylogenetic analysis of the bi-segmented and highly divergent sobemo-like viruses showed a distinctive evolutionary trajectory of its encoding proteins at the periphery of recently reported invertebrate *Sobelivirales*. The beny-like virus belonged to a cluster of insect-associated beny-like viruse; while the picorna-like virus clustered together with psyllid-associated picorna-like viruses. These results highlight the existence of a complex virome within *G. spegazziniana* and establish the basis for future studies investigating the ecological roles, evolutionary dynamics, and potential biocontrol applications of these viruses in the *G. spegazziniana* –yerba mate eco-systems.

## 1. Introduction

The superfamily *Psylloidea* (psyllids) represents an ecologically diverse but relatively understudied group within the suborder *Sternorrhyncha*, order *Hemiptera* [1, 2]. Members of this group, commonly referred to as psyllids or jumping plant lice, are sap-sucking insects highly specialized in their host plants [1, 2]. Some psyllid species are among the most devastating agricultural pests worldwide, either due to their ability to transmit plant pathogens [3] or the range of phenotypes they induce in host plants through salivary secretions [2].

In Argentina, around 16% of the 470 psyllid species known in the Neotropical region have been reported, and several of them are considered economically important pests [4]. *Gyropsylla spegazziniana* (Lizer & Trelles), commonly known as the yerba mate psyllid or “rulo”, belongs to the family *Aphalaridae* [5], and is one of the major pests of yerba mate (*Ilex paraguariensis* A. St-Hil.). This crop holds substantial regional importance, as its leaves and stems are commonly used to prepare a traditional beverage known as ‘mate’ widely consumed in Argentina, Paraguay, Uruguay, and Brazil [6]. *Gyropsylla spegazziniana* induces the formation of leaf deformations, referred to as “galls” or “rulos”, which serve as protective structures for its eggs and nymphs until adulthood, causing important economic losses in yerba mate production [7]. Although several studies have examined the biology, ecology, morphology, and control of *G. spegazziniana* [7-11], its virome remains uncharacterized. This lack of data limits our understanding of how viral communities influence host population dynamics and resilience to environmental stress. Given that viral infections significantly alter arthropod fitness, behavior, and stress tolerance [12], these interactions can dictate pest outbreaks and the success of intervention strategies [13]. Identifying the composition and diversity of the *G. spegazziniana* virome is therefore a critical step toward developing more effective pest management approaches.

The landscape of viral discovery has been fundamentally transformed by the advent of high-throughput sequencing (HTS), which has enabled the systematic characterization of the “dark matter” within the global virosphere [14]. Unlike traditional culture-dependent or PCR-based methods that require prior knowledge of viral sequences, HTS offers an agnostic approach that allows for the detection of viruses in a host-neutral manner [15]. This technological shift has been particularly impactful in the study of invertebrates, where large-scale meta-transcriptomic studies have redefined the known virosphere by uncovering thousands of novel species and filling significant gaps in RNA virus phylogeny [16]. Recent expansive surveys have further extended this reach into aquatic environments, revealing a staggering and previously hidden diversity of RNA viruses within marine invertebrate ecosystems [17]. Furthermore, the sensitivity of HTS allows for high-resolution, in-depth assessments of the viromes associated with individual species, as demonstrated by the characterization of complex viral communities within the golden orb-weaver spider, Nephila clavipes [18]. Collectively, these advancements have reshaped our understanding of the breadth and evolution of the invertebrate virosphere.

Therefore, we implemented the first HTS study on this invertebrate pest and focused on the characterization of the *G. spegazziniana* virome to dig in into the taxonomic composition and diversity of novel viral lineages, unveiling the hidden viral counterpart shaping the psyllid’s biology. By exploring the virome of *G. spegazziniana*, we aimed to provide valuable insights to foster future studies on the ecological roles of viruses within this significant agricultural pest. Moreover, a glimpse of the *G. spegazziniana* virome contributes to a more comprehensive understanding of insect-virus interactions within agricultural ecosystems, potentially impacting broader ecological and evolutionary studies, valuable for the development of novel, virus-based control strategies. This study introduces a fundamental step towards understanding a crucial aspect of this economically important pest and contributing to a more sustainable future for yerba mate crops. Here, we report the identification of five novel RNA viruses, which is, to our knowledge, the first reported viruses associated with this economically important insect, revealing that the *G. spegazziniana* virome is rich and diverse.

## 2. Material and methods

### 2.1. Psyllid sampling and sample processing

Psyllids were collected in yerba mate plants located in Montecarlo, Misiones, Argentina and were used to establish colonies that were maintained at the EEA INTA-Montecarlo greenhouse facility on yerba mate plants under greenhouse conditions [5]. Adults were collected from the artificial rearing population of the psyllid for high-throughput analysis (HTS).

### 2.2. RNA Extraction and HTS Analysis

RNA was extracted from ten pooled psyllids. Insects were ground using a mortar and pestle in liquid nitrogen, and total RNA was extracted using the SV Total RNA Isolation Sytem (Promega, USA) following manufacturer’s instructions. Resulting RNAs were sent to Macrogen INC (South Korea) for library preparation and HTS sequencing. Library was prepared using the TruSeq Stranded Total RNA Library Prep Kit with Ribo-Zero Gold (Illumina, USA). Ribodepleted RNAs were then sequenced in paired-end 101 bp read format on an Illumina NovaSeq6000 platform.

Resulting paired-ends reads were cleaned using the Trimmomatic v0.40 tool, accessed via http://www.usadellab.org/cms/?page=trimmomatic. Standard parameters were employed, apart from raising the quality required from 20 to 30 (initial ILLUMINACLIP step, sliding window trimming, average quality required = 30). Paired, trimmed reads were assembled de novo using rnaSPAdes with standard parameters on the Galaxy server (https://usegalaxy.org/). Resulting contigs were then submitted to local BLASTX searches (E-value < 1 × 10−5) against the complete NR release of viral protein sequences, which was retrieved from https://www.ncbi.nlm.nih.gov/protein/?term=txid10239[Organism:exp]. The resulting viral sequence hits from each dataset were thoroughly examined. Tentative virus-like contigs were curated, extended, and/or confirmed through iterative mapping of filtered reads. This iterative strategy involved extracting a subset of reads related to the query contig, utilizing the retrieved reads to extend the contig, and repeating the process iteratively using the extended sequence as the new query. The extended and polished transcripts were subsequently re-assembled using the Geneious v8.1.9 alignment tool (Biomatters Ltd., Boston, MA, USA) with high-sensitivity parameters.

### 2.3. Sequence and Phylogenetic Analysis

Open reading frames (ORFs) were predicted using ORFfinder with a minimal ORF length of 150 nt and genetic code 1 (https://www.ncbi.nlm.nih.gov/orffinder/). The functional domains and architecture of translated gene products were determined using InterPro https://www.ebi.ac.uk/interpro/search/sequence-search) and the NCBI Conserved domain database-CDD v3.20 (https://www.ncbi.nlm.nih.gov/Structure/cdd/wrpsb.cgi) with an e-value threshold of 0.01. Additionally, HHPred and HHBlits, as implemented in https://toolkit.tuebingen.mpg.de/#/tools/, were employed to complement the annotation of divergent predicted proteins using hidden Markov models. To assess the taxonomic position of the beny-like virus, the full-length replicase proteins were used, while the full-length polyproteins were used for the picorna-like viruses, and the RdRp protein was used for the sobemo-like viruses. Phylogenetic analysis based on these mentioned proteins were carried out using MAFFT 7.526 (https://mafft.cbrc.jp/alignment/software/) with multiple aa sequence alignments using G-INS-i (sobemo-like) and E-INS-i (picorna-like and beny-like) as the best-fit model, respectively. The aligned aa sequences were used as input to generate phylogenetic trees through the maximum-likelihood method with the FastTree 2.1.11 tool available at http://www.microbesonline.org/fasttree/. Local support values were calculated with the Shimodaira-Hasegawa test (SH) and 1000 tree resamples. The sequences of proteins of relevant viral families related to those identified viruses were used as outgroup in the phylogenetic trees.

### 2.4. Nucleic Acid Extraction, RT-PCR Testing, and Sanger Sequencing

Psyllids were collected in yerba mate fields and total RNA was extracted from 10 pooled insects using a modified CTAB protocol (Doyle & Doyle, 1987). Reverse transcription was performed using 5 μL of the total nucleic acid extract in a 20 μL reaction mixture that contained 5× first-strand buffer (Promega, Madison, WI, USA), 1 μL 10 mM dNTP, 1 µl 250ng/ul random hexamers,, and 0.8 μL M-MLV reverse transcriptase (Promega). Before the reverse transcription reaction, the RNA template was incubated at 70 °C for 5 min, then the reverse transcription mix was added. The profile used consisted of incubation at 37 °C for 60 min and reverse transcriptase deactivation at 70 °C for 10 min. All polymerase chain reactions (PCR) were accomplished by Pegasus Taq polymerase (Productos Bio-Lógicos SA, Argentina) in a 25 μL reaction mixture that contained 2.5 μL 10× Buffer, 1 μL 10 mM dNTP,1 μL 10 μM of each forward primer and reverse primer, 2.4 units of Pegasus Taq Polymerase, 15.9 μL free RNAse/DNAse water and 2 μL cDNA template. The PCR profile consisted of denaturing at 94 °C for 5 min, and 40 cycles of 94 °C for 40 s, 53-60 °C for 45 s (depending on the melting temperature of primers used), 72 °C for 1 min, followed by a final extension for 10 min at 72 °C. Testing for the presence of each one of the five identified viruses was conducted using those primers listed in the Supplementary Table S1; which were designed according to the assembled sequences for each one of the five viruses. For Sanger sequencing, PCR fragments were purified with with Gel/PCR DNA Fragments Extraction Kit (Geneaid Biotech Ltd, Taiwan), and submitted for sequencing to Macrogen, Inc. (South Korea).

## Results and Discussion

### 3.1 Viruses associated with G. spegazziniana

Insects are the most abundant group of animals on earth [19] . These animals have a rich and diverse virome which is being uncovered in the later years by the use of HTS [16, 20-24]; however, only a small portion of the insect viromes diversity has been unraveled so far [25-26]. The viromes of just four psyllids species (*Bactericera cockerelli, Diaphorina citri; Leuronota fagarae* and *Trioza erytreae*), were characterized so far and several viruses were identified [27-35]. Moreover, the data mining of picornavirales from publicly available insect RNA-seqs datasets resulted in the identification of two novel viruses associated to two distinct psyllid species [36]; highlighting the potential richness of the virome associated to psyllids. To date, around 4,000 psyllid species have been described and classified [1], each exhibiting distinct life histories and host exploitation strategies [2]. This shows that the viruses linked to psyllids is highly limited, poorly misrepresents the diversity of these invertebrates, and even important agronomic pests as the yerba mate psyllid virome are still unexplored. Therefore, in this study, we explored the virome of *G. spegazziniana*, which resulted in the identification of five novel highly divergent RNA viruses, including a beny-like virus, a picorna-like virus, and three sobemo-like viruses, which were tentatively named Gyropsylla spegazziniana beny-like virus 1 (GSBlV1), Gyropsylla spegazziniana picorna-like virus 1 (GSPlV1), and Gyropsylla spegazziniana sobemo-like virus 1 (GSSlV1), Gyropsylla spegazziniana sobemo-like virus 2 (GSSlV2), Gyropsylla spegazziniana sobemo-like virus 3 (GSSlV3). The first exploration of the virome associated to *G. spegazziniana* highlights how rich and diverse is the virome associated with this highly important yerba mate pest. We assembled the full-length coding region for all viruses, and we also detected all these viruses in field collected psyllids suggesting that these viruses are circulating in *G. spegazziniana* populations. Revealing the virus diversity in this pest is crucial to further exploit the identified viruses as specific and targeted tools for the biological control of insect populations.

### 3.2. Molecular and phylogenetic characterization of the newly identified yerba mate psyllid Beny-like virus

The *Benyviridae* is a family with members that have positive-sense, single stranded RNA viruses with only one genus, the *Benyvirus*, recognized by the ICTV [37]. Benyviruses are plant-infecting viruses with a multipartite genome [37-38]. Nevertheless, metagenomic studies as well as the metatrancriptomic analysis of public data deposited in the NCBI, resulted in the discovery of many benyvirus-related sequences, that resemble to newly unassigned Benyviridae members, that likely infect a wide range of non-plant hosts, such as insects and fungi [39-45]. Furthermore, the insect-associated beny-like virus genomes are monopartite which is contrasting to the multipartite genomes displayed by the plant associated counterparts [39]. Here, a novel beny-like virus was identified and named as Gyropsylla spegazziniana beny-like virus 1 (GSBlV1). The positive-sense, single stranded RNA GSBlV1 sequence (GenBank accession number PZ145639) is 7024 nucleotides (nt) long and has two ORFs encoding a non-structural polyprotein of 1945 amino acids (aa) and a putative structural protein of 274 aa (Figure 1A) (Table 1). This genomic organization is similar to that reported for other insect-associated beny-like viruses [39-45], as well as for Pistacia ribo-like virus (PiRbLV; MT334604; Mohammadi et al., unpublished). GSBlV1 non-structural polyprotein showed the highest BlastP similarity with that one encoded by PiRvLV, with 57.92% identity (Table 1); therefore, based on genetic distance, GSBlV1 is a new virus distantly related to other NCBI database entries of insect viruses. Three conserved domains, Alphavirus-like methyltransferase (MT) domain, the Viral_helicase1 (Hel), and the ps-ssRNAv_RdRp-like (pol), were identified in the polyprotein (Figure 1A) at aa positions 60–282, 591–757 and 1584–1881, respectively. Similar conserved domains were found in those non-structural polyproteins encoded by related beny-like viruses [39-40,42-44]. Unlike its plant-associated counterparts [38, 44], insect-associated beny-like virus replicase, like those encoded by GSBlV1 and its related viruses [39-40, 42-43], do not have any papain-like protease domain in their sequences. GSBlV1 structural protein showed the highest BlastP similarity with the one encoded by PiRbLV, with 55.35% identity (Table 1). This protein encoded by the GSBlV1 ORF2 is likely a coat protein, since the TMV_coat super family conserved domain was identified at aa position 123–274. A similar conserved domain was also identified in the ORF2-encoded proteins of other insect-associated beny-like viruses [39, 43].

**Figure 1.**
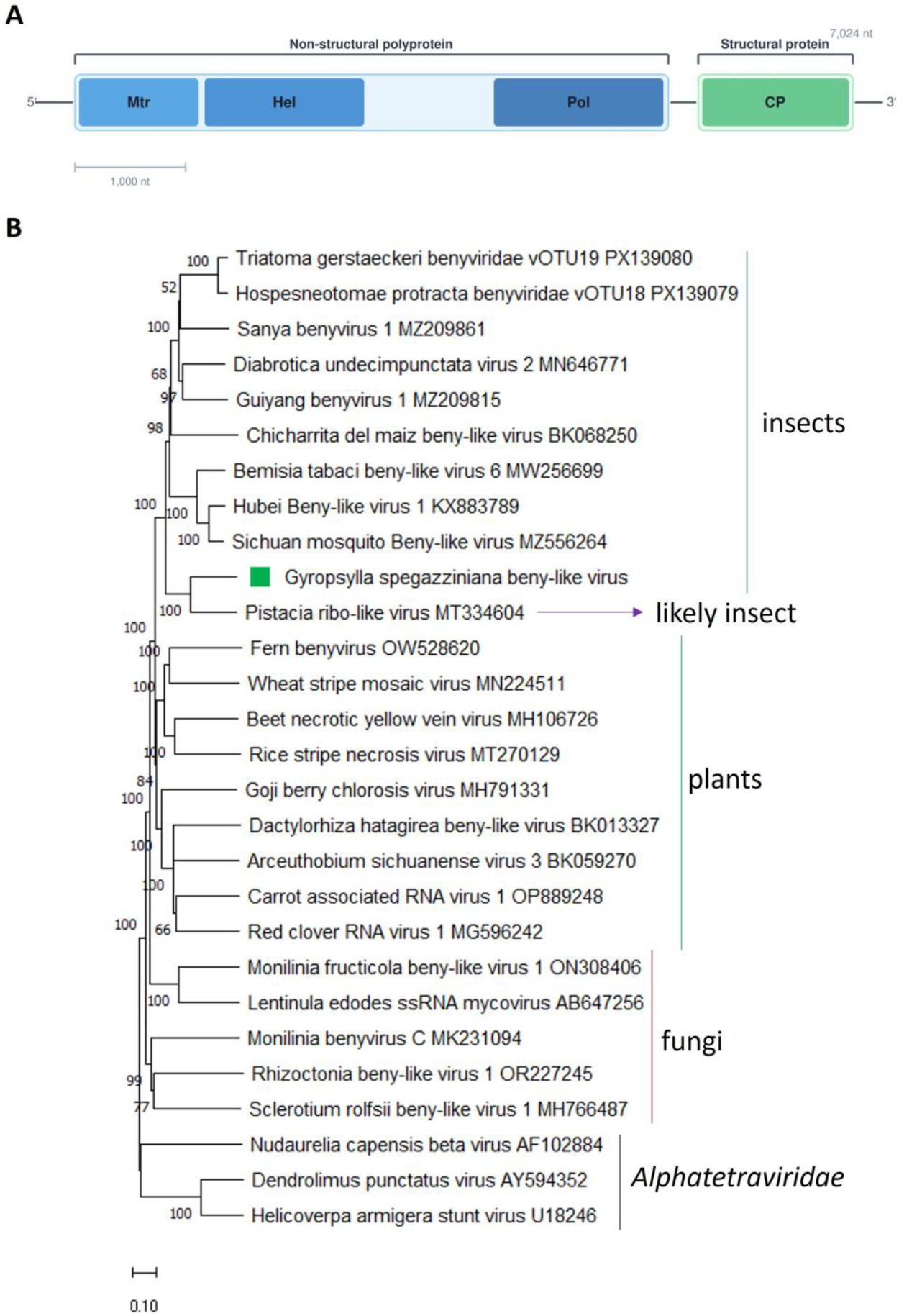
(A) Schematic representation of the genome organization of Gyropsylla spegazziniana beny-like Virus 1. Mtr, methyltransferase; Hel, helicase; Pol, RNA polymerase; CP, capsid protein. (B) Maximum likelihood phylogenetic trees reconstructed using the replicase protein sequence of Gyropsylla spegazziniana beny-like virus 1 and of representative beny-like viruses. Bootstrap values above 50% are shown (1000 replicates). Gyropsylla spegazziniana beny-like 1 virus is indicated with a green square. The scale bar shows the substitution per site.

Phylogenetic analysis of the non-structural polyprotein (replicase) amino acid sequence placed GSBlV1 in a clade containing several insect-associated beny-like viruses, as well as one associated to Pistacia, where GSBlV1 was clustered together with the PiRbLV. Thus, it is highly likely that PiRbLV is associated with an unknown insect, whose RNA that was co-purified along with Pistacia RNA. The insect-associated clade of beny-like viruses, as well as PiRbLV is separated from the clades composed of fungi-associated beny-like viruses and plant-associated ones (Figure 1B). Thus, we support the proposal made by Debat et al., [39] to create a new genus, to be named as “Insebenyvirus” within the family *Benyviridae*, to accommodate the insect-associated beny-like viruses, as well as PiRbLV. Based on the phylogenetic insights and the observed genetic distance of the newly identified viruses, we tentatively propose an aa sequence identity of 80% in the Replicase or/and CP as threshold for species demarcation in this newly proposed genus.

To our knowledge, GSBlV1 is the first reported beny-like virus associated with psyllids.

### 3.3. Molecular and phylogenetic characterization of the newly identified yerba mate psyllid Picorna-like virus

The order *Picornavirales* is comprised of nine families, and many of its members are associated to arthropod hosts [46]. This order contain viruses with a positive-strand ssRNA genome [47], and the viral RNAs of all members of the *Picornavirales* are polyadenylated but can display distinct genome organizations, e.g., monopartite or bipartite genomes which encode one to five ORFs [46]. Here, a novel picorna-like virus was identified and named as Gyropsylla spegazziniana picorna-like virus 1 (GSPlV1) (GenBank accession number PZ145640). GSPlV1 has a positive-sense 9804 nt RNA genome encoding a single 2922 aa polyprotein (Fig. 2A, Table 1).

**Figure 2.**
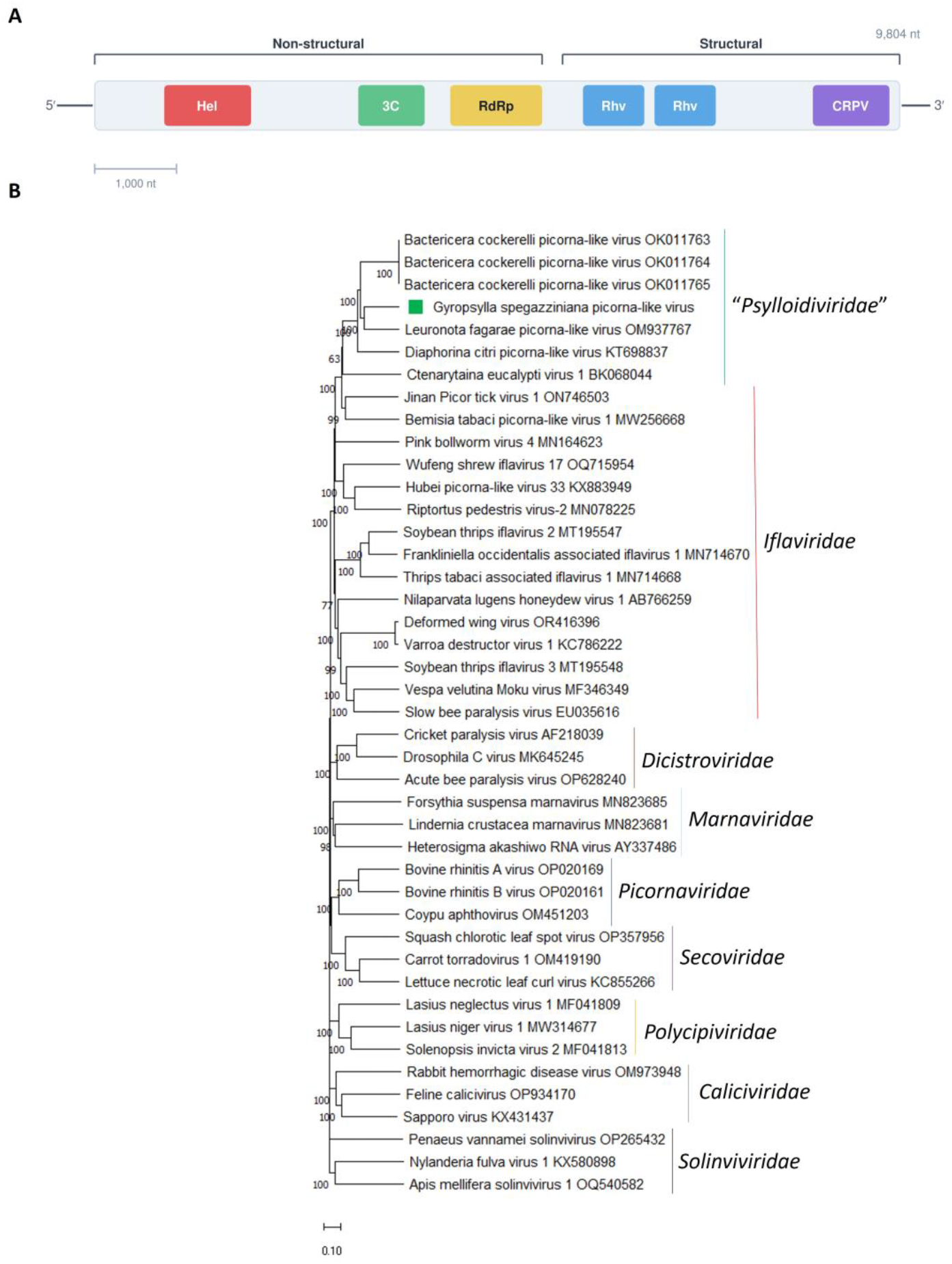
(A) Schematic representation of the genome organization of Gyropsylla spegazziniana picorna-like virus 1. Hel, helicase; 3C, 3C-like protease; RdRp, RNA-directed RNA polymerase; Rhv, rhinovirus-like capsid protein; CRPV, cricket paralysis virus-like capsid (B) Maximum likelihood phylogenetic trees reconstructed using the polyprotein protein sequence of Gyropsylla spegazziniana picorna-like virus 1 and of representative picornavirids. Bootstrap values above 50% are shown (1000 replicates). Gyropsylla spegazziniana picorna-like virus 1 is indicated with a green square. The scale bar shows the substitution per site.

GSPlV1 polyprotein showed the highest BlastP similarity with that one encoded by the psyllid associated Leuronota fagarae picorna-like virus (LfPLV), with 57.31% identity (Table 1). Genomic organization of GSPlV1 is similar to that one reported for other psyllid-associated picorna-like viruses [28, 30, 34, 36], where the non-structural domains are located at the N-terminal region of the polyprotein, while the structural domains are located downstream in the C-terminal region (Figure 2B). A helicase (RNA_Helicase, aa 666–840), a Picornavirales 3C/3Clike protease (3C-Pro, aa 1241–1448), and a RNAdirected RNA polymerase (RdRp, aa 1477–1985) conserved domains were identified in the N-terminal region of the GSPlV1polyprotein (Figure 2A); while two rhinovirus-like capsid domains (Rhv-like, aa 2003–2244 and aa 2267–2512) and a cricket paralysis virus–like capsid domain (CRPV_capsid superfamily, aa 2685–2914) were identified in the C-terminal region of the GSPlV1 polyprotein (Figure 2A).

Iflaviruses use a type IV (dicistrovirus-like) IRES within the 5’ UTR for cap-independent translation [48]. Secondary structure prediction of the GSPlV1 5’UTR employing the RNAfold web server showed a highly compact, multi-domain architecture containing several hallmark IRES motifs, which are known to be critical for ribosomal recruitment [30], suggesting that this region likely functions as an IRES. Like it was reported for Diaphorina citri picorna-like virus [30], and LfPLV [34] the putative GSPlV1 IRES is distinct to the type IV IRES found in iflaviruses, supporting Stuehler et al., [34] and Du et al., [30] suggestion that this distinct architecture may represent a novel variant or type.

Conserved domains are typical of picornaviruses [30, 46] supporting the placement of GSPlV1 within the order *Picornavirales*. Phylogenetic analysis based on the polyprotein shows that GSPlV1 clustered within the psyllid-associated clade (Figure 2B) supporting Du et al., [30] assumption that psyllid associated viruses share a recent common ancestor, which had an inverted genomic architecture regarding that one displayed for iflaviruses. GSPlV1 and other psyllid-associated viruses cluster close to members of the *Iflaviridae* family, but forming a distinct clade (Figure 2B). Moreover, GSPlV1 possesses several genomic characteristics, also described for the other psyllid associated viruses [28, 30, 34, 36] that undoubtedly differentiate it from *Iflaviruses* and indicating that these viruses should not be classified into any established family, most particularly: (1) different genome organization, the non-structural proteins are located in the N-terminal region, while structural proteins are located in the C-terminal region of the viral genome, an arrangement opposite to that of all known iflaviruses; (2) a structurally divergent 5’ UTR IRES element; and (3) less than 40% polyprotein sequence identity with any recognized iflaviruses. Therefore, GSPlV1 should be included as a new member of the recently proposed new family, tentatively named “*Psylloidiviridae*”, within the proposed new genus, tentatively named “*Psylloidivirus*” [30]. The identification of GSPlV1 as well as a novel psyllid associated virus [36] strongly support the formal recognition of this newly proposed taxa by the ICTV.

Several picornaviruses have been reported to be harmful to the host insects leading to morphological deformities, behavioral change, and death [21].Thus, further studies should explore the potential pathogenicity to their insect host of this newly picorna-like virus associated to yerba mate psyllids for the potential development as a tool for biocontrol psyllids population in yerba mate.

### 3.4. Molecular and phylogenetic characterization of the newly identified yerba mate psyllid Sobemo-like virus

*Solemoviridae* is a recently established family of positive-strand RNA viruses whose recognized members are associated to plants [49]. Usually, solemovirids have unsegmented genome with four to ten ORFs, which code for a viral suppressor of RNA silencing, a polyprotein (which contains a serine protease and an RdRp conserved domains), and a CP [49]. Nevertheless, recently, sobemo-like viruses with two segments were identified in arthropods such as crustaceans [50], myriapods [16], and insects [33, 51-54]. Here, three sobemo -like viruses were identified and named as Gyropsylla spegazziniana sobemo-like virus 1 (GSSlV1) (GenBank accession number PZ145641-PZ145642), Gyropsylla spegazziniana sobemo-like virus 2 (GSSlV2) (GenBank accession number PZ145643-PZ145644) and Gyropsylla spegazziniana sobemo-like virus 3 (GSSlV3) (GenBank accession number PZ145645-PZ145646) (Table 1), which increases the group of sobemo-like viruses with bi-segmented genomes associated to non-plant hosts. A sobemo-like virus was previously identified in the psyllids *Bactericera cockerelli* [29] and *Trioza eritreae* [33]. Thus, *G. spegazziniana* is likely the third psyllid host where sobemo-like viruses associated to these insects have been identified.

The GSSlV1, GSSlV2 and GSSlV3 genomes comprised two segments of single-stranded, positive-sense RNA (Figure 3A-C, Table 1). RNA 1 is composed of 3482, 3418 and 3391 nt, respectively; while RNA 2 comprises 1506, 1694 and 1706 nt, respectively (Table 1).

**Figure 3.**
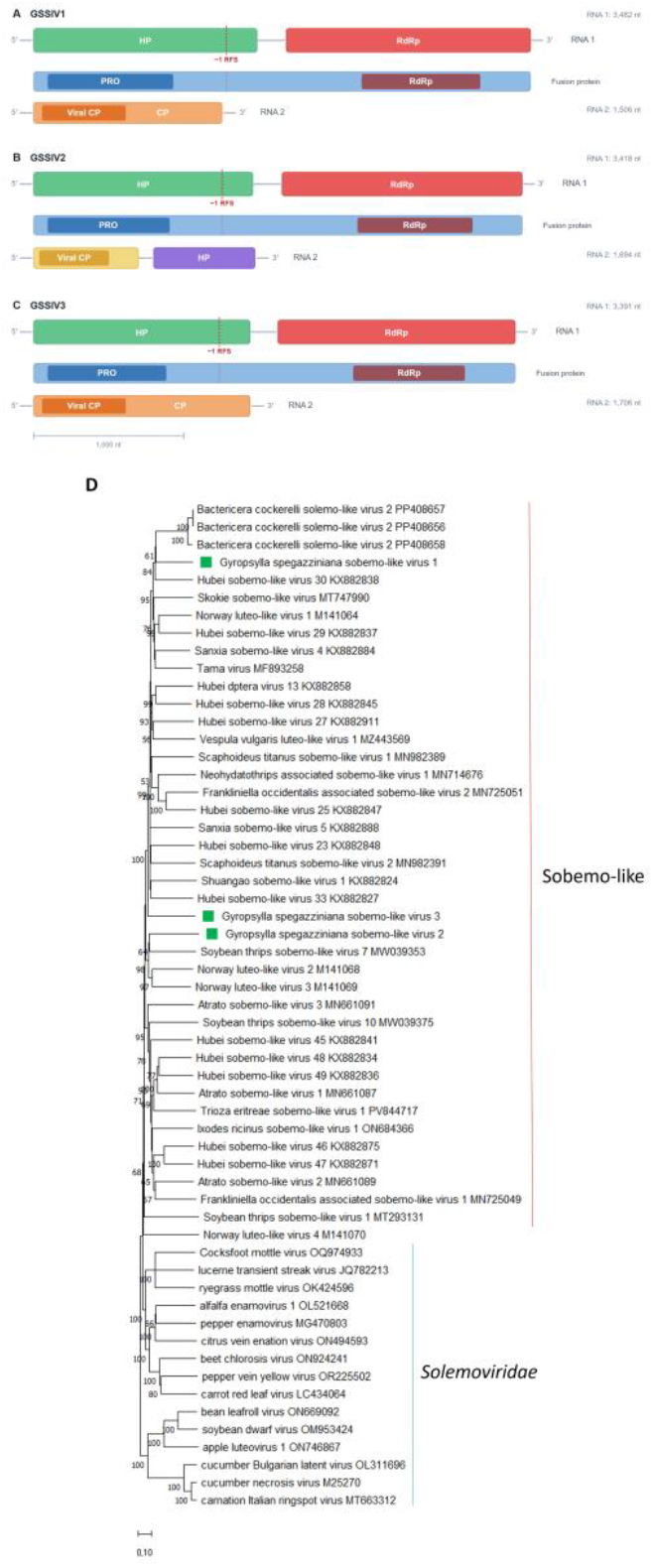
(A) Schematic representation of the genome organization of Gyropsylla spegazziniana sobemo-like virus 1 (B) Schematic representation of the genome organization of Gyropsylla spegazziniana sobemo-like virus 2 (C) Schematic representation of the genome organization of and Gyropsylla spegazziniana sobemo-like virus 3. Pro, Trypsin-like serin protease; RdRp, RNA-directed RNA polymerase; Viral CP, viral_coat (D) Maximum likelihood phylogenetic trees reconstructed using the RdRp protein sequence of Gyropsylla spegazziniana sobemo-like virus 1, Gyropsylla spegazziniana sobemo-like virus 2 and Gyropsylla spegazziniana sobemo-like virus 3 and of representative sobemo-like viruses and plant-associated solemovirids. Bootstrap values above 50% are shown (1000 replicates). Gyropsylla spegazziniana sobemo-like virus 1, Gyropsylla spegazziniana sobemo-like virus 2 and Gyropsylla spegazziniana sobemo-like virus 3 are indicated with green squares. The scale bar shows the substitution per site.

GSSlV1, GSSlV2 and GSSlV3 segment 1 have two ORFs named HP and RdRP. ORF HP encodes a hypothetical protein (HP), and an RdRP is encoded by the other ORF and is expressed as a fusion polyprotein through a −1 ribosomal frameshift mechanism (Figure 3A-C). GSSlV1 HP has 713 aa and shows the highest BlastP similarity with Bactericera cockerelli solemo-like virus 2 with 36.65% identity; while GSSlV2 HP has 672 aa and shows the highest BlastP similarity with Culex inatomii luteo-like virus with 28.42% identity, and GSSlV3 HP has 593 aa and shows the highest BlastP similarity with Moscow lipoptena solemo-like virus with 29.54% identity. A Trypsin-like serin proteases (PRO) conserved domain was identified in GSSlV1, GSSlV2 and GSSlV3 HPs (Figure 3A-C) at aa positions 241–404, 185-344, and 213-363, respectively. GSSlV1 RdRp has 396 aa and shows the highest BlastP similarity with Bactericera cockerelli solemo-like virus 2 with 59.39% identity; while GSSlV2 RdRp has 345 aa and shows the highest BlastP similarity with Tartas insect-associated virus with 41.03% identity, and GSSlV3 RdRp has 375 aa and shows the highest BlastP similarity with Sanxia sobemo-like virus 4 with 46.53% identity. A viral RNA-dependent RNA polymerases (RdRp) conserved domain was identified in GSSlV1, GSSlV2 and GSSlV3 RdRps (Figure 3A-C) at aa positions 59–303, 49-248, and 37-257, respectively.

The genomic organization of segment 1 of GSSlV1, GSSlV2 and GSSlV3 as well as in those sobemo-like viruses associated to arthropods is highly similar to that one reported for the 5′ terminal half region of the solemovirids (the ORF encoding the viral suppressor of RNA silencing protein is not present, since plants are not their hosts), as the first ORF encodes a polyprotein with a serine protease domain; while the RdRp is translated via −1 programmed ribosomal frameshift (−1 PRF) from the next ORF [55].

The segment 2 of GSSlV1 and GSSlV3 encode a CP (Figure 3A and C) with a size of 416 aa and 435 aa, respectively. GSSlV1 CP shows the highest BlastP similarity with Scaphoideus titanus sobemo-like virus 1 with 36.41% identity; while GSSlV3 shows the highest BlastP similarity with Neohydatothrips associated sobemo-like virus 1 35.44%. On the other hand, the segment 2 of GSSlV2 encodes two proteins (Figure 3B), a CP with a size of 219 aa, and a HP of 248aa. The CP shows the highest BlastP similarity with Chihuahua culicoides solemo-like virus 1 with 33.62% identity, while the HP shows the highest BlastP similarity with Tryoza eritreae sobemo-like virus with 30.19% identity. A viral_coat (Viral CP) conserved domain was identified in GSSlV1, GSSlV2 and GSSlV3 CPs (Figure 3A-C) at aa positions 59-229, 65-234 and 21-216, respectively; on the other hand, any conserved domain was identified in the HP encoded by the segment 2 of GSSlV2. Interestingly, any segment 2 was reported for the psyllid-associated Bactericera cockerelli solemo-like virus 2 [29]. Thus, it is tempting to speculate that authors did not search that segment in their data. Therefore, we analyzed the SRA28090463 associated to the bioproject PRJNA1080349 [29] which resulted in the identification of the putative segment 2 of the Bactericera cockerelli solemo-like virus 2 with 1617 nt, that encodes a 415aa CP, which shares a 47.8% identity to the one encoded by GSSlV1. This finding supports that all arthropod-associated sobemo-like viruses have two segments, despite that only one segment was reported for many of the already reported sobemo-like viruses associated to arthropods. Moreover, GSSlV1, GSSlV2 and GSSlV3, as well as the other identified sobemo-like viruses CPs, also share sequence identities with noda-like viruses and permutotetra-like viruses, which has been linked with the CP being encoded by subgenomic RNAs or genomic segments in these viruses, so it is likely that horizontal gene transfer could have been the consequence of mispackaging during co-infection of the same host [55].

Phylogenetic analysis based on the RdRp sequences, placed GSSlV1, GSSlV2 and GSSlV3 within the clade containing unclassified solemovirids identified from a variety of arthropods; where GSSlV1 clustered with the psyllid-associated Bactericera cockerelli solemo-like virus 2 and Hubei sobemo-like virus 30, GSSlV2 clustered together with soybean thrips sobemo-like virus 7, and GSSlV3 is placed in a monophyletic position (Figure 3D).

GSSlV1, GSSlV2 and GSSlV3 share very low identity in their encoded proteins and are located in distinct clades, indicating that these three viruses likely have distinct evolutionary trajectories.

Taking into account the phylogenetic relationships as well as genomic organization of solemovirids associated to non-plant hosts, a novel family, with several genera to be further assigned, should be created within the order *Sobelivirales* to accommodate the solemovirids that have arthropods as their hosts.

Furthermore, this distinct genomic arrangement of the solemovirids associated to non-plant hosts should led to further research into its replicative mechanisms and evolutionary origins.

### 3.5. GSBlV1, GSPlV1, GSSlV1, GSSlV2 and GSSlV3 can be found in field-collected psyllids

Psyllids were field-collected in yerba mate crops, and tested for the presence of the five identified viruses using RT-PCR with those primers listed in Supplementary Table S1. All five viruses were successfully amplified from the field-collected samples, RNA 1 segment of GSSlV1, GSSlV2 and GSSlV3 was targeted, showing the expected size of the PCR bands when tested with the appropriate primers for each virus (Supplementary Figure S1). These amplified fragments were submitted for Sanger sequencing which resulted in nucleotide sequences more than 99% identical to those derived from the HTS assembled sequences for the five viruses. Further studies could be conducted using the designed primers in order to assess the infection rate of each one of the five identified viruses across *G. spegazziniana* populations in distinct sampling sites as well to assess the viral dynamics on the yerba mate psyllids populations to determine the potential role of these five psyllid viruses in controlling psyllid numbers in yerba mate crops. Similar studies in other psyllids are being conducted [29, 56] which will be useful to find out viral dynamics in psyllids populations and the role of the viruses in controlling the psyllids populations.

#### Concluding remarks

Our transcriptome analysis of the yerba mate psyllid, led to the identification of five novel viruses which have broaden the knowledge about the psyllid virosphere, shedding light on the diversity and phylogenetic relationships of viruses infecting this economically important pests. Thus, metagenomics and metatranscriptomics have huge potential for uncovering even greater viral diversity within psyllid species. The detection of these five identified viruses in field-collected insect samples, suggest that these viruses are widespread among the yerba mate psyllid populations. Our research unlocks thrilling ways for exploring the functional roles of these newly discovered viruses within *G. spegazziniana*. Conducting virus–host interaction studies and functional assays will be key to get essential knowledge into how these viruses influence their host’s physiology, behavior, and susceptibility to environmental factors.

Understanding these interactions could led to the development of novel and sustainable pest-control strategies, potentially targeting specific viral processes or manipulating virus–host interactions to interfere with the yerba mate psyllid biology. Moreover, the knowledge of the virome (insect viral community) of insect pests is key to develop bio-insecticides based on viruses; therefore, the identification of the yerba mate psyllid virome would be useful to develop novel and sustainable approaches based on virocontrol to manage this agronomically important pest.

## Author Contributions

**YSC**: conceptualization, investigation, formal analysis, writing- review and editing. **AB**: investigation, formal analysis, writing- review and editing. **VN**: investigation, writing- review and editing. **HD**: investigation, formal analysis, data curation, writing- review and editing **MES**: investigation, funding acquisition, writing- review and editing. **NEB**: investigation, formal analysis, data curation, conceptualization, funding acquisition, writing- original draft. All authors have read and agreed to the published version of the manuscript.

## Institutional Review Board Statement

This article does not contain any studies with human participants or animals performed by any of the authors.

## Informed Consent Statement

Not applicable.

## Data Availability Statement

The sequences were deposited in GenBank with accession numbers PZ145639-PZ145646.

## Figure Legends

**Supplementary Figure S1.**
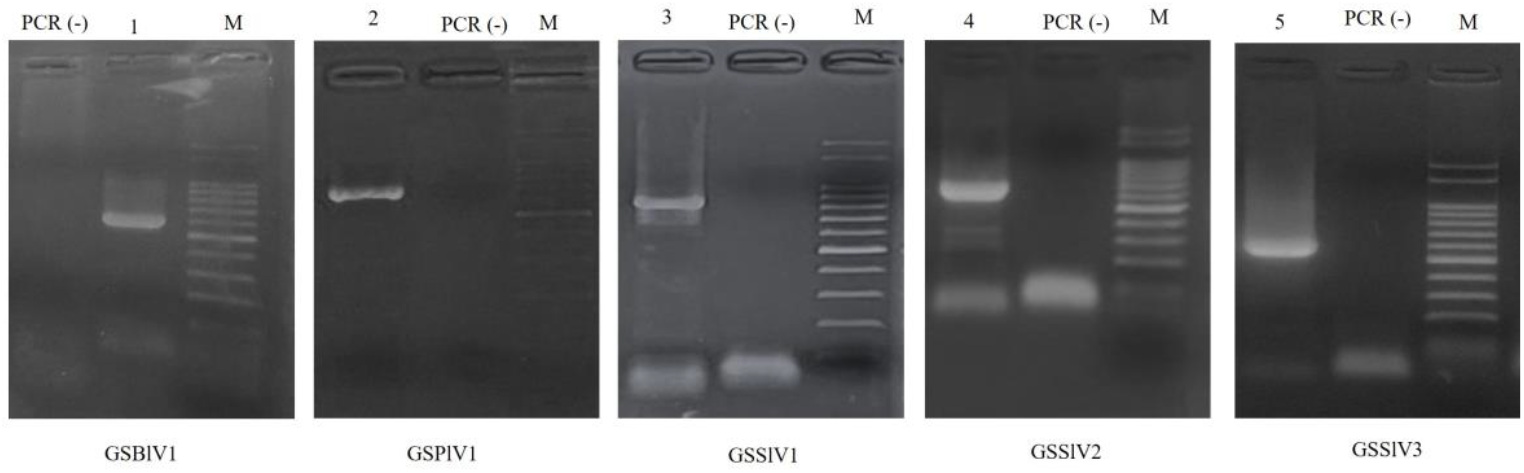
Testing the five viruses by endpoint RT-PCR in a bulked sample of ten field-collected yerba mate psyllids; M: molecular marker, 100 bp DNA ladder (PB-L Productos Bio-Lógicos, Argentina (100, 200, 300, 400, 500, 600, 700, 800, 900, 1000, 1500 and 2000 pb)). PCR (-): PCR negative control. 1) Gyropsylla spegazziniana beny-like virus 1 (GSBlV1) (703pb), 2) Gyropsylla spegazziniana picorna-like virus 1 (GSPlV1) (566pb), 3) Gyropsylla spegazziniana sobemo-like virus 1 (GSSlV1) (890pb), 4) Gyropsylla spegazziniana sobemo-like virus 2 (GSSlV2) (759pb), and 5) Gyropsylla spegazziniana sobemo-like virus 3 (GSSlV3) (793pb).

**Supplementary Table S1.**
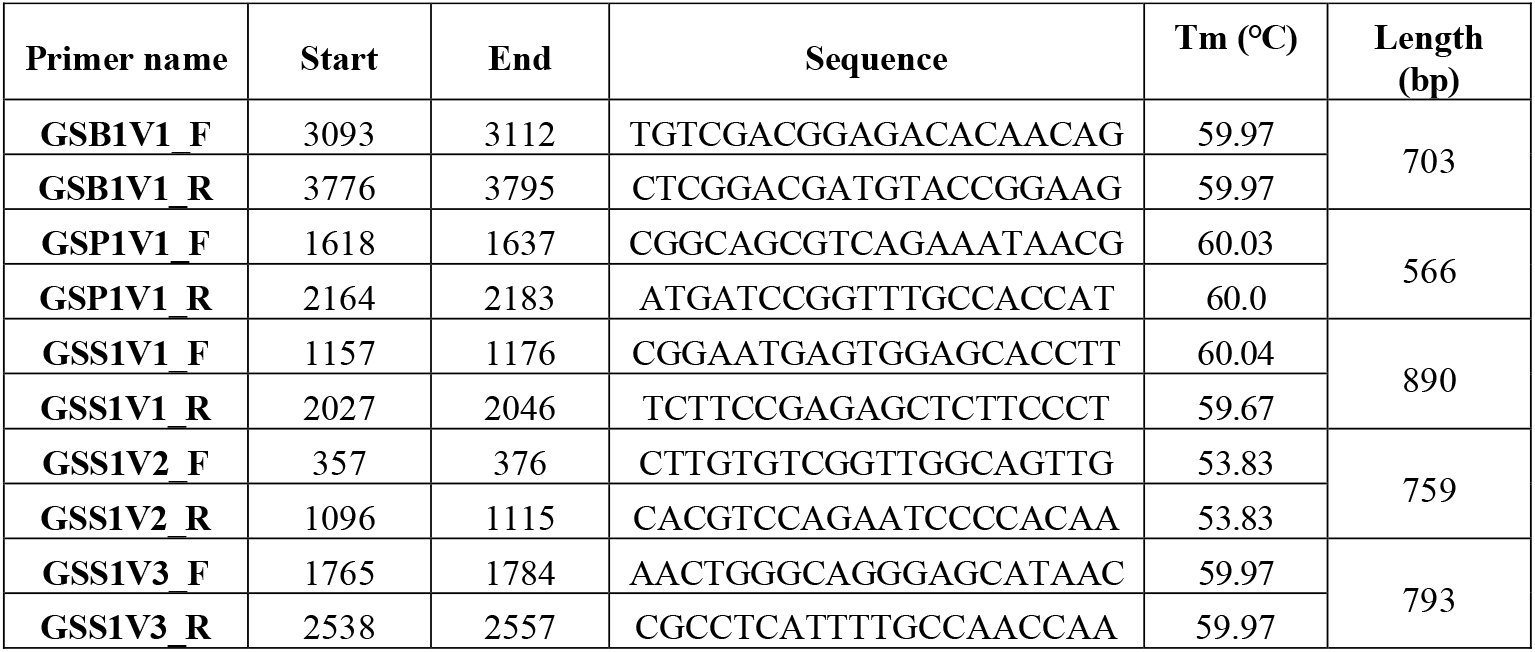
List of primer pairs used in this study. The sequences for all primer pairs used for RT-PCR testing of each virus (GSB1V1, GSP1V1, GSS1V1, GSS1V2 and GSS1V3) are given in the table, with start and end positions based on the corresponding contig assembled from HTS data as escribed in main text, Tm for each primer as calculated using Primer-BLAST (NCBI) with default parameters, and length of predicted PCR product.

